# Do honey bee species differ in the odometer used for the waggle dance?

**DOI:** 10.1101/2020.12.17.423277

**Authors:** Ebi Antony George, Neethu Thulasi, Patrick L. Kohl, Sachin Suresh, Benjamin Rutschmann, Axel Brockmann

## Abstract

Honey bees estimate distances to food sources using image motion experienced on the flight path and they use this measure to tune the waggle phase duration in their dance communication. Most studies on the relationship between experienced optic flow and the dance-related odometer are based on experiments with *Apis mellifera* foragers trained into a small tunnel with black and white patterns which allowed quantifiable changes in the optic flow. In this study we determined the calibration curves for foragers of the two Asian honey bee species, *A. florea* and *A. cerana*, in two different natural environments with clear differences in the vegetation conditions and hence visual contrast. We found that the dense vegetation condition (with higher contrast) elicited a more rapid increase in the waggle phase duration with distance than the sparse vegetation in *A. florea* but not in *A. cerana*. Visual contrast did not affect the perception of the food reward, measured as the number of dance circuits produced per distance, in both species. Our findings suggest that contrast sensitivity of the waggle dance odometer, or other aspects of flight behaviour, might vary among honey bee species.

## Introduction

Honey bees, like other flying insects, mainly use image motion (optic flow) which is dependent on the environment to estimate their flight distances (Esch and Burns, 1996; Lecoeur et al., 2019; Srinivasan et al., 1996; Srinivasan, 2011). Studies in *Apis mellifera* established that it is this optic flow driven odometer, and not, for example, time of flight or energy consumption, that is used to tune the waggle phase duration, the distance signal of the dance communication (Dacke and Srinivasan, 2008; Esch et al., 2001; Srinivasan et al., 2000). The odometer is based on contrast information in the green spectral channel and even very low levels of contrast are sufficient to detect and process image motion information (Chittka and Tautz, 2003; Si et al., 2003). Most of these findings are based on studies in which foragers were trained into a small tunnel which allows a controlled manipulation of the visual environment (Esch et al., 2001; Srinivasan et al., 2000).

Experiments exploring the variation of waggle run duration in natural habitats with apparent differences in optic flow information are rare (Esch et al., 2001; Tautz et al., 2004). Tautz and colleagues (2004), for example, compared waggle run durations for feeder locations on land and water. In line with their expectation, the slope for the relation between feeder distance and waggle run duration (“calibration curve”) was higher on land than on water. However, water is an extreme environment with respect to honey bee foraging, and the authors reported that no recruits visited the feeder in the middle of the lake. Thus, our knowledge of whether and to what extent honey bee colony calibration curves of honey bee colonies vary between natural environments which differ in their vegetation density is scarce (Collett, 2000). In this study, we present a comparison of calibration curves of the same colonies between a dense and a sparse vegetation density environment for two different Asian honey bee species, *Apis florea* and *Apis cerana*.

*Apis florea*, the open nesting red dwarf honey bee species, is phylogenetically most distant from the cavity nesting species, like *A. cerana* and *A. mellifera* (Raffiudin and Crozier, 2007; Smith, 2020). *A. florea* markedly differs from the two very similar cavity nesting species in size, nesting environment and in the variety of signals produced in the waggle dance (Dyer, 2002; I’Anson Price and Grüter, 2015). The waggle dance in the ancestral *A. florea* occurs on the horizontal surface at the top of the comb, the crown, and is directed towards the food source (Dyer, 1985). In the cavity nesting *A. cerana*, foragers perform dances on vertical combs (Lindauer, 1956; von Frisch, 1967). In spite of these differences, the two species show, strong similarities in the relationship between the waggle phase duration and foraging range, the change in dance precision with distance and the dance follower behaviour (Beekman et al., 2015; George et al., 2020; Kohl et al., 2020; Sen Sarma et al., 2004). *A. florea* and tropical *A. cerana* have similarly “steep” calibration curves which increase with distance more rapidly than the temperate *A. mellifera* (Kohl et al., 2020). Given that *A. florea* and *A. cerana* apparently have a more similarly tuned waggle dance odometer while being phylogenetically most distant, they are well suited to explore possible differences or the dance odometer among honey bee species.

We studied the effect of two different natural environments, with different vegetation conditions and hence visual contrast, on four aspects of the waggle dance behaviour; the waggle phase duration, the total circuit duration, the return phase duration and the number of dance circuits performed per dance in *A. florea* and *A. cerana*. Crucially, we tested foragers from the same colony in both the dense and sparse vegetation conditions by shifting the whole colony between the conditions in both species. We were interested in exploring the effect of optic flow information on the waggle dance behaviour in these Asian *Apis* species and compare this with results from *A. mellifera*.

## Materials and Methods

### Colony preparation and experimental location

The experiments were performed with a natural *Apis florea* colony located on the National Centre for Biological Sciences, Bangalore campus and an *A. cerana* colony bought from a commercial beekeeper. In both species, the colony was first shifted to the dense vegetation condition with high visual contrast in the Botanical Garden of the University of Agricultural Sciences, Gandhi Krishi Vignana Kendra, Bengaluru (Fig. 1A, Fig. S1). After experiments in the Garden, the colony was shifted to an open field in the same campus with sparse vegetation and hence low visual contrast (Fig. 1B, Fig. S1). In the case of *A. florea*, the colony was placed in a box which allowed video recordings of the crown area of the colony, which is where foragers dance (Dyer, 1985). In the case of *A. cerana*, the colony was placed in an observation box with the individual frames horizontally next to each other. The wall of the box nearest the entrance was made of glass, allowing recordings of the dance floor area. The experiments in the botanical garden are described elsewhere (Kohl et al., 2020), although the dances for this manuscript were analysed separately.

**Figure 1:**
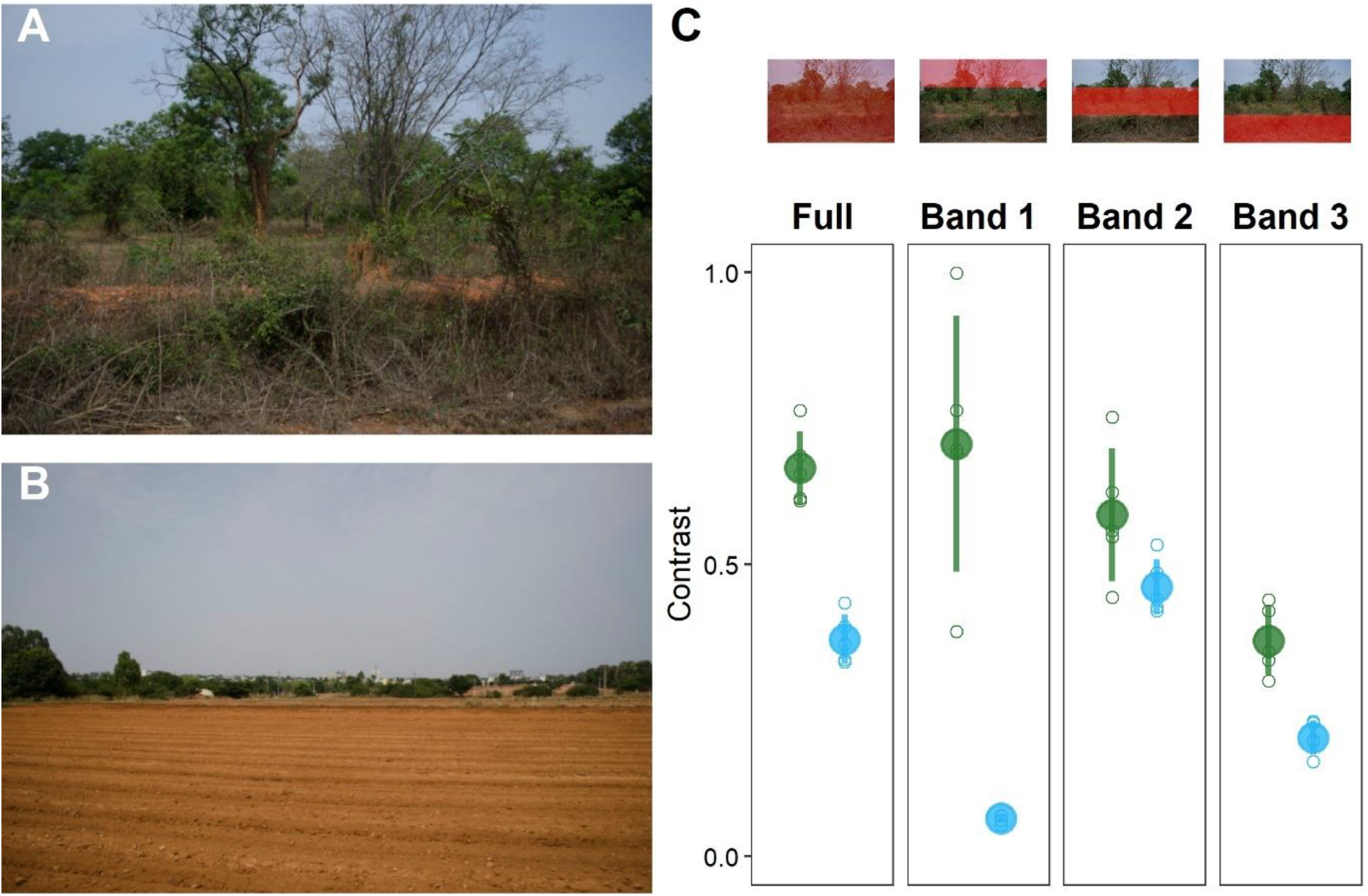
Representative images of A) Dense and B) Sparse vegetation conditions, taken in the direction away from the hive location, towards the feeder location. C) The amount of contrast present in the 5 images from each condition. The coloured circles and error bars correspond to the mean and standard deviation of the two conditions (green: Dense vegetation, blue: Sparse vegetation) and the grey circles correspond to the contrast obtained from the individual images. The Dense vegetation condition had greater contrast than the Sparse vegetation condition in all comparisons, although this was not significant in Band 2.

### Experimental protocol

The same experimental protocol was followed for the experiments in both the vegetation conditions and in both the species. The colony was shifted into the location on the evening of day 0, to ensure that most foragers have returned to the colony before the shift. After a day’s break, the colony training was initiated on day 2. Foragers from the colony were trained to an artificial feeder filled with sucrose solution placed next to the colony initially. The sucrose concentration of the feeder was adjusted between 1 to 1.5 M, depending on the number of foragers visiting the food source. Once 10-20 foragers were coming to the feeder on their own, the feeder was shifted in steps of 5-10 m to 25 m. At 25 m, the foragers were individually marked (Uni POSCA Paint Markers, Uni Mitsubishi Pencil, UK).

The feeder was then shifted in small steps to 100 m and the foragers coming to this distance were observed in the hive. Their dance activity was recorded using a Sony HDR-CX240 at 50 frames per second. After an hour of recordings, the feeder was shifted to the next distance of 200 m and the dances by the foragers active at this distance were recorded for another hour. This protocol was repeated for distances of 300 m, 400m and 500 m over a period of 5 days. Since feeder training had to be done from the beginning for both the vegetation conditions, we could not ensure that the same individual foragers were active at our feeder in the two conditions.

### Contrast Analysis

To confirm the difference in the visual contrast between the dense and the sparse vegetation conditions, we analysed the amount of contrast in images from the two conditions following the protocol in Tautz et al., 2004. We obtained 5 images each, at a resolution of 4928 x 3264 pixels, from the two conditions. The images were obtained at distances of 100 m, 200 m, 300 m, 400 m, and 500 m facing in the direction away from the hive towards the feeder (Fig. 1A and 1B). We then obtained the average of the intensity of the red, green, and blue channels for each pixel. The contrast was then calculated as the standard deviation of these average pixel intensities divided by the mean of the average pixel intensities. We calculated the contrast using 4 different sections of the image: the full image, a horizontal band covering the top third of the image (Band 1), a horizontal band covering the middle third of the image (Band 2) and a horizontal band covering the bottom third of the image (Band 3). We then compared the level of contrast in each of these sections between the two vegetation conditions. We compared between contrast in the whole images as well as contrast in the different bands to check whether the overall difference in contrast was reflected in arbitrary smaller sections of the image.

We used a linear model with the contrast value as the response and the vegetation condition as the predictor and found that overall, there was a strong difference in the contrast between images from the Dense and Sparse vegetation conditions (Fig. 1C, difference estimate between Dense and Sparse = 0.293, t = 8.59, p < 0.0001). To compare the contrast levels within the three horizontal bands, we first compared two linear models; one with the contrast as the response, and the condition and the band identity as the predictor, and another with the same response and predictor variables, but with an interaction term between the predictors. We found that the model with interaction was significantly better at explaining the data (F = 18.375, p < 0.0001). We then checked the model with interactions and found that the Dense vegetation condition had higher contrast than the Sparse vegetation condition, although the difference was not significant in Band 2 (Fig. 1C, Band 1: difference estimate between Dense and Sparse = 0.642, t = 9.56, p < 0.0001; Band 2: difference = 0.124, t = 1.84, p = 0.078;

### Video analysis

We followed established protocols for analysing the waggle dance activity (Seeley, 1995). The honey bee waggle dance consists of a waggle phase, which is very short at shorter distances, and a return phase (Gardner et al., 2008). The first frame in which a bee moved its abdomen dorsoventrally or laterally while starting a forward motion was considered as the start of the waggle phase and the last frame in which the bee stopped moving its abdomen before turning away from a straight line path was considered as the end of the waggle phase. The return phase was the time period between the end of one waggle phase and the start of the next. The circuit duration for one dance circuit was the summation of the waggle phase duration and the subsequent return phase duration. The number of dance circuits corresponds to the number of waggle phases in the dance. We analysed a total of 34 (dances per distance: mean ± standard deviation = 6.8 ± 2.39) and 36 (mean ± standard deviation = 7.2 ± 1.79) dances respectively in the Dense and Sparse vegetation conditions for *A. florea* and a total of 50 dances each (10 dance per distance) for both conditions in *A. cerana* (Table S1).

### Statistical Analysis

#### Effect of visual contrast on waggle dance behaviour

Our statistical analysis of the effect of visual contrast on the waggle dance behaviour in both species focused on 4 waggle dance parameters: the waggle phase duration, the total circuit duration, the return phase duration and the number of dance circuits. We present results from the analysis on the waggle phase duration and the number of dance circuits in the main text, and the results of the analysis on the total circuit duration and the return phase duration in the supplementary text. The analysis was done separately for the data on *A. florea* and *A. cerana.* We used mixed effects models to determine the effect of distance and vegetation condition on each of the parameters while accounting for differences amongst bees in the slope of the relationship between the parameter of interest and distance. The models had the respective parameter as the response variable, an interaction between the distance (a continuous variable) and the visual contrast condition (a categorical variable of two levels) as the predictor, and Bee ID as the random effect on the slope.

In the case of the three parameters which were continuous in nature (the waggle phase duration, the total circuit duration and the return phase duration), the analysis was done on the mean duration of all the runs in each dance, and not on the individual runs themselves. We first fit linear mixed effects models (LMMs) for all these three response parameters. However, in most cases, model assumptions were not validated due to the non-linear distribution of the data (Fig. 2A and 3A), and as a result, we do not make inferences from the LMMs. Instead, we fit a non-linear mixed effects model (NLMM) with the logarithmic regression as the non-linear function. The logarithmic regression is of the form:

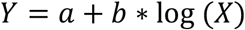

**Figure 2:**
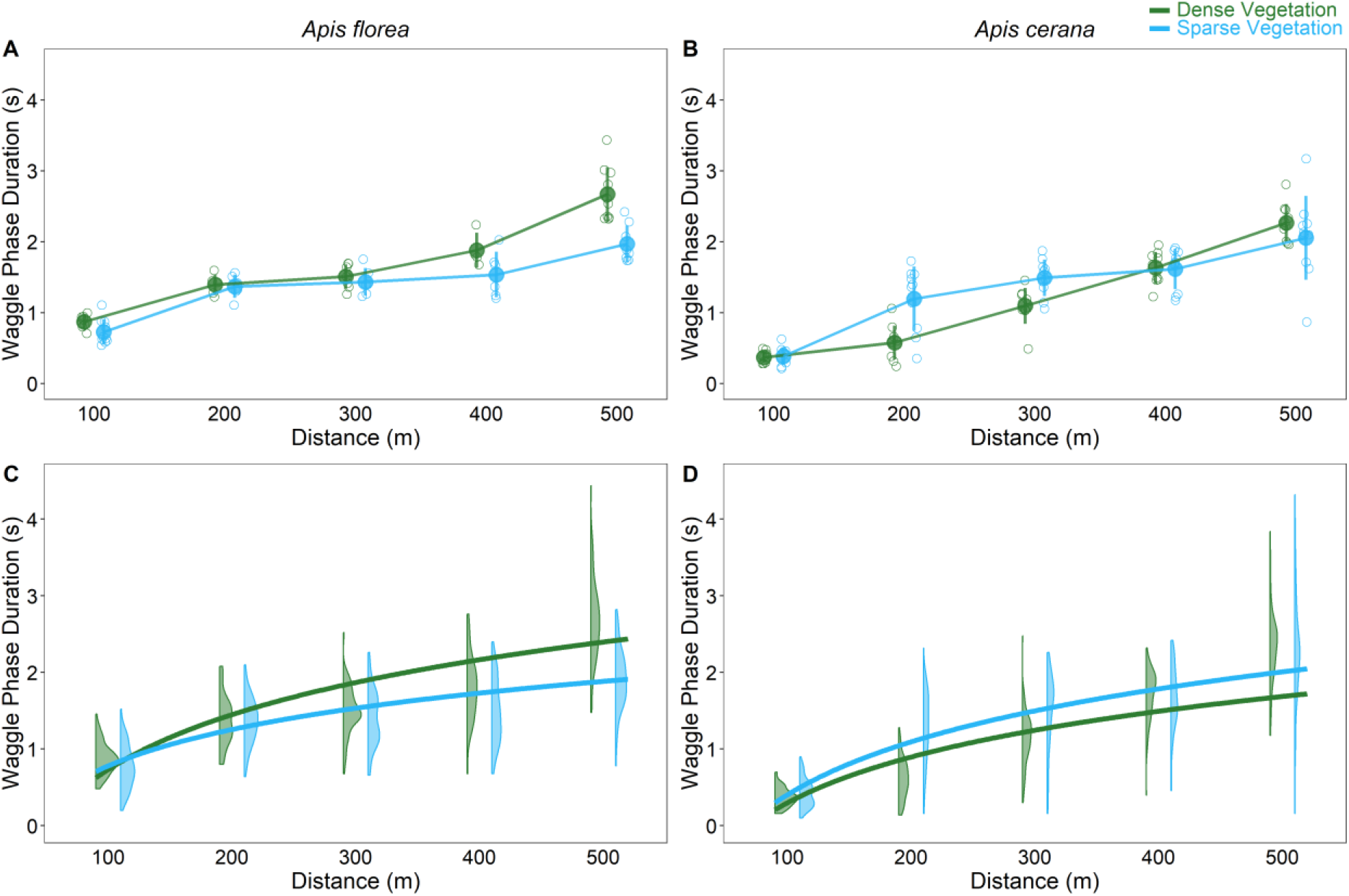
The waggle phase duration at different distances in the two vegetation conditions in *A. florea* (A, C) and *A. cerana* (B, D). A and B) The mean waggle phase duration for individual dances (open circles), along with the mean of all the dances and the standard deviation (closed circles with error bars) for *A. florea* and *A. cerana*, respectively. C and D) The predicted fits obtained from the non-linear mixed effects logarithmic regression model overlaid on top of the distribution of individual runs at each distance for *A. florea* and *A. cerana*, respectively. Circles, fitted lines and the distributions are coloured based on the vegetation condition, with green for Dense vegetation and blue for Sparse vegetation.

**Figure 3:**
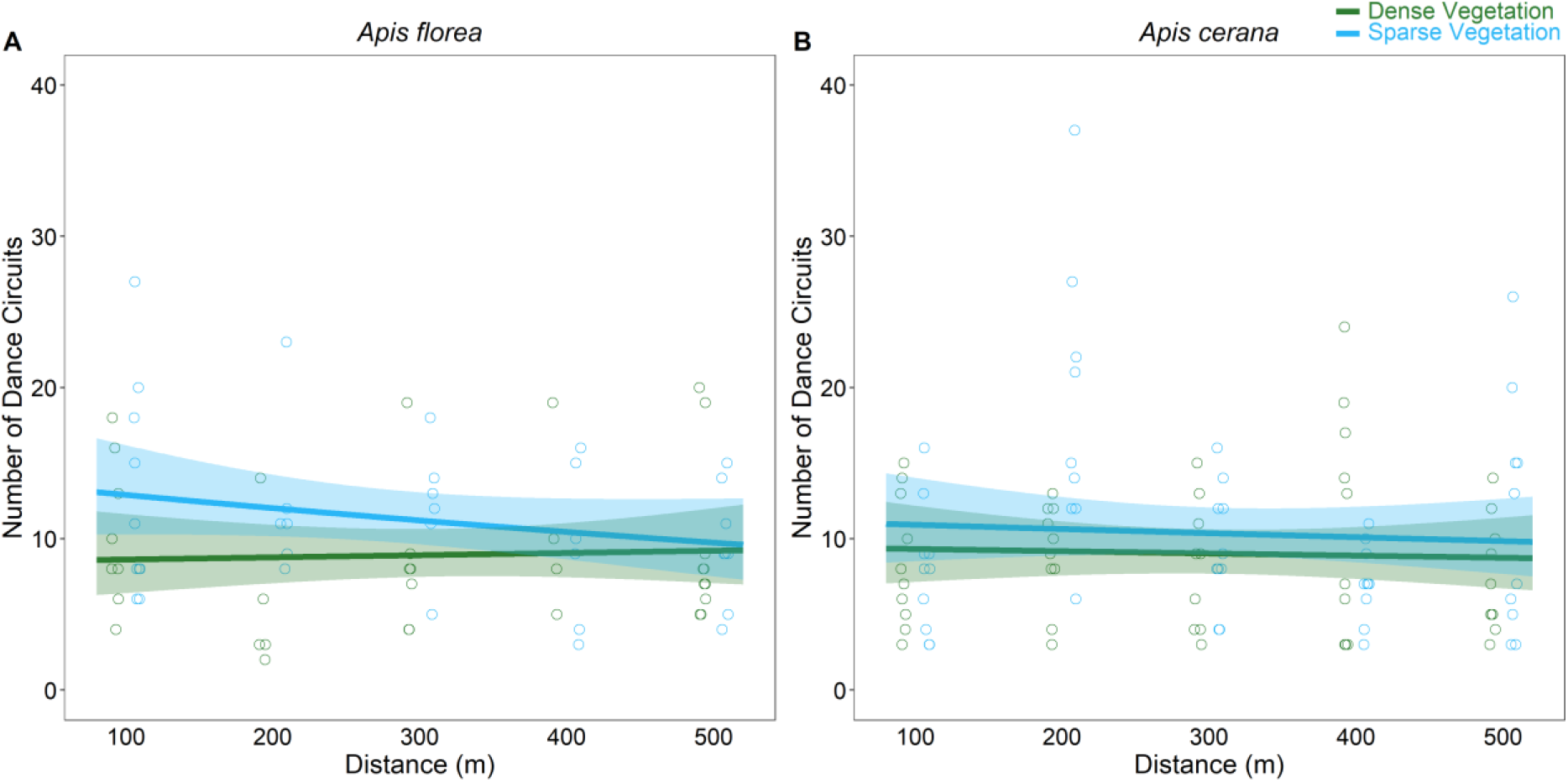
The number of dance circuits at different distances in the two vegetation conditions in A) *A. florea* and B) *A. cerana*. The open circles represent the number of dance circuits in each individual dance, with the lines and shaded region representing the predictions and confidence interval of the predictions at the fixed effects level obtained from the negative binomial mixed-effects model. The circles, lines and shaded region are coloured based on the vegetation condition, with green for Dense vegetation and blue for Sparse vegetation.

The NLMMs were used to estimate and compare the value of *a* and *b* for both vegetation conditions. Since *a* and *b* are analogous to the intercept and slope in a linear regression, for simplicity’s sake we refer to them as the intercept and slope henceforth. We are primarily interested in quantifying the effect of the vegetation conditions on the slope *b* in the different dance parameters. Further details regarding the fitting of the non-linear models, including obtaining the starting values, are provided in the supplementary text. We verified the model assumptions of homoscedasticity and normality of the residuals for NLMMs, and they were validated in all except one case. In the case of the waggle phase duration in *A. cerana*, there were signs of heteroscedasticity in the NLMM. To account for this, we provided a power variance function structure in the model (see Supplementary text). The number of dance circuits is a discrete parameter, and we fit a negative binomial mixed effects model to account for the discreteness as well as the overdispersion in the data.

Further, we performed three other confirmatory analysis and we have provided their details in the supplementary information accompanying this manuscript. In the first analysis, for *A. florea*, we fit LMMs for the waggle phase duration, as model assumptions were not violated. We then compared the results obtained from the LMM and the NLMM described above. In the second analysis, we focussed on the change in variation in the waggle phase durations with distance under the two visual contrast conditions in both species. We were interested in looking at whether visual contrast can modulate the variation shown in the waggle dance, in addition to its effect on the waggle phase duration itself, and if this can account for some of the non-linearity seen in the data we obtained. In the third analysis, we focused on the effect of potential dances with intermediate waggle phase durations on the results seen in our main analysis. On location of a new food sources, most foragers in *A. mellifera* take a few dances to update their waggle duration. In these intermediate dances, the waggle phase duration shown by the forager is in between the waggle phase duration for the old and the new food, source location (Chatterjee et al., 2019). To account for this, we removed dances immediately after the shift to the new location from our data and performed the analysis again.

All the statistical analysis was performed in R 4.0.1 (R Core Team, 2018), using the RStudio IDE (RStudio Team, 2016). We used the aomisc (Onofri, 2020) and nlme package (Pinheiro et al., 2020) to fit the LMMs and the NLMMs and the glmmTMB package (Brooks et al., 2017) to fit the negative binomial mixed effects models. We used ggplot2 (Wickham, 2016), gghalves (Tiedemann, 2020) and cowplot (Claus O. Wilke, 2018) packages to visualise the model fit and the data. Additionally, we used pillow (Clark, 2020), numpy (Oliphant, 2006) and pandas (McKinney and others, 2010) packages in python (Van Rossum and Drake, 2009) for extracting the contrast values from photographs.

## Results and Discussion

The principal finding of our study is that the foragers of *A. florea* and *A. cerana* colonies responded differently to the two visual environments as indicated by the dance-distance calibration curves. All calibration curves showed an increase with feeder distance independent of environment. However, only for the *A. florea* foragers the slope of the calibration curve in the visually more cluttered environment was significantly higher by 1.5-fold compared to that in the open field (Fig. 2, S2 and Table S2, slope: Dense = 1034.51, Sparse = 685.69; difference estimate = −348.82, confidence interval = −581.21 - -116.43, t = - 2.921, p = 0.005). For *A. cerana* foragers the calibration curves for the two environments did not significantly differ (Fig. 2 and Table S3, slope: Dense = 862.88, Sparse = 997.61; difference estimate = 134.73, confidence interval = −62.13 – 331.59, t = 1.336, p = 0.186). The circuit duration also followed the same pattern in *A. florea* and *A. cerana* (*A. florea* - Fig. S3 and Table S2, slope: Dense = 2470.37, Sparse = 1268.26; difference estimate = −1202.11, confidence interval = −2132.23 - -271.99, t = −2.515, p = 0.015; *A. cerana* - Fig. S3 and Table S3, slope: Dense = 1768.24, Sparse = 1357.31, difference estimate = −410.93, confidence interval = −968.59 – 146.73, t = −1.438, p = 0.155). These results suggest that the estimation of flight distances and subsequent dance information in natural environments might be more complex and plastic than predicted by flight responses in spatially restricted tunnels with controlled visual black and white patterns.

Some of these tunnel experiments showed that the dance odometer in *A. mellifera* is driven by contrast in the green spectral channel (Chittka and Tautz, 2003) and that a large change in contrast between vertical stripes (from 20% to 92%) leads to a 1.5-fold change in the waggle, run duration for the same tunnel distance (Si et al., 2003). Unfortunately, there is only one study so far that tried to estimate the effect of optic flow on the waggle dance in two visually distinct natural environments, over land and over water (Tautz et al., 2004). A change in contrast from around 9% (over water) to around 20% (over land) led to a 4-fold increase in the slope of the calibration curve. So, both types of experiments clearly indicated that contrast can affect distance estimation and waggle phase duration. However, the relation between contrast differences in the landscape and perceived optic flow is unclear. For example, Si et al. (2003) suggested that their results favoured a threshold model and concluded that the odometer is robust against a large range of contrast variation, beyond a minimum contrast level. However, their results are not that as clear as they stated, as there were still changes in the waggle phase duration beyond this threshold. On the other hand, the contrast differences in the field and tunnel experiments were calculated in different ways. In the field experiments, the contrast was calculated from sections of images which recaptured the assumed visual scene experienced by the bee whereas the contrast calculation in the tunnel experiments was based on the whole flight path experienced by the bee. Obviously more experiments are needed to clarify the effect of contrast variation on the flight and dance odometer.

Nonetheless, based on previous studies, we expected to observe significant differences in the slopes between the dense and sparse vegetation in both species. Thus, the question arises why we did not find differences in the slopes of the calibration curves for *A. cerana* but did so in *A. florea*. Since contrast has an effect on the flight odometer, differences in contrast sensitivity among honey bee species could affect the calculation of flight distances and waggle run durations. Interestingly, a recent tunnel experiment with *A. cerana* and *A. mellifera* foragers reported clear differences in the flight responses to artificial black and white patterns between the foragers of both species (Chakravarthi et al., 2018). The authors of the study concluded that the behavioral differences suggest strong differences in the spatial resolution and contrast sensitivity used for flight manoeuvres, which were not expected based on the morphological similarities of the eyes of *A.mellifera* and *A. cerana* (Kelber and Somanathan, 2019; Streinzer et al., 2013). Furthermore, honey bee species might differ in their flight responses, e.g., flight height, in different environments which can in turn affect the odometer (Collett et al., 2006).

In contrast to the results regarding the flight and dance odometer, the parts of the dance that are related to the perceived food reward – number of dance circuits and duration of the return phase - showed the expected correlation (Barron et al., 2007; George and Brockmann, 2019; Hrncir et al., 2011; Łopuch and Tofilski, 2020; Seeley, 1989, 1986; Seeley et al., 2000; Shafir and Barron, 2010; von Frisch, 1967). In both species, the number of dance circuits was not affected by distance or the vegetation conditions (*A. florea* - Fig. 3 and Table S2, slope: Dense = 0.0002, Sparse = −0.0007; difference estimate = −0.0009, confidence interval = - 0.0023 – 0.0006, z = −1.165, p = 0.244; *A. cerana* - Fig. 3 and Table S3, slope: Dense = - 0.0002, Sparse = −0.0003; difference estimate = −0.0001, confidence interval = −0.0015 – 0.0014, z = −0.134, p = 0.893), whereas the return phase duration slightly increased with distance but was not affected by vegetation conditions (*A. florea* - Fig. S4 and Table S2, slope: Dense = 1407.33, Sparse = 592.00; difference estimate = −815.33, confidence interval = −1732.49 – 101.84, t = −1.729, p = 0.089; *A. cerana* - Fig. S4 and Table S3, slope: Dense = 699.94, Sparse = 408.35; difference estimate = −291.59, confidence interval = −787.37 – 204.20, t = −1.147, p = 0.255). Similar to previous distance training experiments in honey bees, we used a high sugar concentration to keep the foragers motivated to dance and recruit throughout the experiment (Seeley, 1995; von Frisch, 1967). This likely led to a ceiling effect, and that prevented a decline in the number of dance circuits with distance (Seeley, 1994). Interestingly, we still could observe a significant increase in the duration of the return phase with distance. Seeley at al. (2000) demonstrated an effect of the reward value on the return phase by increasing the sugar concentration of a feeder at one distance: the higher the sugar concentration the faster the return phase. This response makes the dance appear more intense or “lively” as Lindauer called it (Boch, 1956; Lindauer, 1948). An increase in the distance of a food source with the same energetic value should lead to a lowered perception of the reward value (due to higher energy costs associated with flight) and an accompanying increase in the return phase duration, consistent with our results. Alternatively, the increase in return phase duration could also be linked to the increase in waggle phase duration with distance, which necessitates a longer return phase duration to get back to the starting point of the waggle phase and complete the waggle circuit (Heran, 1956). Interestingly, the results on the return phase duration differ from the observations on *A. mellifera* by Tautz et al., (2004), who reported that the slope of the return phase duration with distance over water (i.e., low visual contrast) was higher than that over land. However, the authors of the study note that they did not find any recruits at the feeder when the food source was over water (Tautz et al., 2004). The presence of the food source at an unnatural location (over water) for honey bee foragers in that study could explain the unexpected results they obtained regarding the return phase duration.

To conclude, we are convinced that our small study raises new questions regarding the working of the flight and dance odometer in honey bees. The effect of contrast on the odometer is still unclear. However, the odometer might be more robust against ecologically realistic variation of the visual terrain than a comparison of dance durations over water and land in *A. mellifera* had suggested. In addition, our results indicate that there might be differences in the effect of contrast on the odometer amongst honey bee species. Also, honey bee species may differ in aspects of their flight behaviour, e.g., flight height, that affect the perception of contrast and thus the odometer. More experiments are needed focussing on the relationship between species-specific differences in the visual system and flight behaviour that drive the odometer used for the waggle dance.

## Acknowledgements

We would like to thank the Department of Apiculture, GKVK Bangalore for allowing us to perform experiments on their premises. We would also like to thank Mahesh Kumar MH and Sruthi Unnikrishnan for help during the experiments.

## Author Contributions

Conceptualization: E.A.G., P.L.K., A.B.; Methodology: E.A.G., P.L.K., A.B.; Formal

analysis: E.A.G., N.T., S.S.; Investigation: E.A.G., N.T., P.L.K., B.R.; Writing - original, draft: E.A.G., A.B.; Writing - review & editing: E.A.G., N.T., P.L.K., S.S., B.R., A.B.; Supervision: A.B.; Funding acquisition: A.B.

