## Supplementary Information for "Do honey bee species differ in the odometer used for the waggle dance?"

N.T.: 0000-0003-4702-8900

P. L. K.: 0000-0001-9278-978X

S.S.:  0000-0002-5245-7008

B. R.: 0000-0001-6589-6408

A.B.: 0000-0003-0201-9656

* corresponding author:

Ebi Antony George

In our experiments, waggle phase duration nor circuit duration showed a simple linear relationship with distance in both species. Further, we found that in *A. cerana*, the mean waggle phase durations, and the spread of the accompanying distribution, for the lower distances were higher in the sparse vegetation conditions as opposed to the dense vegetation conditions (e.g., 200 m distribution, Fig. 2B). We performed three confirmatory analysis to verify our dataset.

#### LMM for Waggle Phase duration in *A. florea*

In the first analysis, we built a linear mixed effects model (LMM) to compare the slope of the calibration curve (change in waggle phase duration with distance) in the two vegetation conditions in *A. florea*. We could do this for *A. florea*, but not for *A. cerana* as the homoscedasticity model assumption was violated in the case of the latter. We then compared the results obtained from the non-linear mixed effects model (NLMM) described in the main text with the results of this LMM.

We found that the LMM also gave the same results as the NLMM. The waggle phase duration increased with distance, but this increase was significantly higher in the Dense vegetation condition as compared to the Sparse vegetation condition (Fig. S2; slope: Dense = 4.31, Sparse = 2.86; difference estimate = -1.45 confidence interval = -2.34 - -0.55, t = -3.16, p = 0.003).

#### Variation Analysis

One possibility for the difference in variation in the two vegetation conditions is that visual contrast can modulate the variation of the waggle phase durations, in addition to its effect on the mean duration. In our second analysis we compared the variation in the waggle phase duration with distance and vegetation conditions in both *A. florea* and *A. cerana*. We obtained the standard deviation of the waggle phase duration for each dance by each individual forager and then built linear mixed-effects models with the standard deviation values as the response, an interaction between distance and visual contrast condition as the predictor and bee ID as the random effect. Further, we also calculated the coefficient of variation (standard deviation/mean) for the waggle phase durations for each dance, and performed the same statistical analysis

**Results**

*A. florea*

The standard deviation of the waggle phase increased with distance, but the slope was higher for the dense vegetation condition as opposed to the sparse vegetation condition (Fig S5 and Table S4, slope: high = 0.530, low = 0.275; difference estimate = -0.255, confidence interval = -0.508 - -0.001, t = -2.014, p = 0.049). The coefficient of variation did not show a significant increase with distance in either vegetation condition (Fig S5 and Table S4, slope: high = -0.00016, low = -0.00033; difference estimate = -0.00016, confidence interval = -0.00039 - -0.00006, t = -1.457, p = 0.151).

*A. cerana*

The standard deviation of the waggle phase increased with distance, but there was no significant difference in the slopes between the two vegetation conditions (Fig S5 and Table S5, slope: high = 0.759, low = 0.777; difference estimate = 0.018, confidence interval = -0.374 – 0.410, t = 0.092, p = 0.927). The coefficient of variation showed a decrease with distance in both vegetation conditions, but the slopes were not different between the two conditions (Fig S5 and Table S5, slope: high = -0.00027, low = -0.00023; difference estimate = 0.00003, confidence interval = -0.00028 - 0.00035, t = 0.217, p = 0.829).

**Discussion**

We found interesting differences between the two species in both measures of variation in the waggle phase duration, the standard deviation and coefficient of variation. As expected with an increase in the mean, the standard deviation increased with distance in both conditions for both species (Beekman et al., 2015). However, while the slope of this increase was higher in the high vegetation conditions for *A. florea*, the increase was the same in both vegetation conditions for *A. cerana*. In the case of the coefficient of variation, there was no significant change with distance in both vegetation conditions in *A. florea*. But in *A. cerana*, the coefficient of variation slightly decreased with distance, although there were no differences between the two vegetation conditions.

The coefficient of variation (CV) is a better measure compared to the standard deviation, as it takes into account the increase in variation associated with an increase in the mean value of a distribution. The similarity in the slopes for the CV in both vegetation conditions in both species indicates that visual contrast does not affect the variation in the waggle phase duration in either species.

However, inferences from these results must be made cautiously as we are limited in our sampling for understanding the variation in the waggle phase per distance (range: 4 -10 data points) and the number of colony replicates (1 colony per species). Further studies would be needed to observe whether: a) visual contrast can modulate the standard deviation of waggle phase durations, as seen in our data on *A. florea* and b) the coefficient of variation decreases with distance as seen in our data on *A. cerana*.

#### Effect of Intermediate Dances

On finding food at a new location, not all honey bee foragers immediately update their waggle phase duration to reflect the new location of the food source (Chatterjee et al., 2019). Most foragers can take up to 3 waggle dances to show the changed waggle phase duration. In the dances before the change-point, these foragers show waggle phase durations which are intermediate between the durations for the old location and the new location. In our experiments, we could not ensure that all dances that were analysed were not intermediate dances. This is because we had a short time window in which to perform the experiments. *Apis florea* and *A. cerana* are known to abscond relatively easily as compared to *A. mellifera*. In addition, the shifting of the colonies to two new locations within weeks increased the likelihood of the colonies absconding. However, to estimate the effect that these intermediate dances have on the dance slopes in the high and low vegetation conditions we did a secondary analysis. In our data for *A. cerana*, we calculated the dance number after the shift for each dance analysed for each individual forager. We then produced a second dataset removing all the dances that would potentially have intermediate waggle phase durations (first to third dance at each distance) and performed the same analysis as in the main manuscript on this dataset. In total, after removing intermediate dances, we had 57 dances (out of the initial 100) in our second dataset.

**Results**

We found that the same distributions fit the data best for the second dataset also. On fitting the appropriate distributions, we obtained estimates of the slopes that were similar to the slopes obtained in our main analysis (Table S6). The pattern of the slopes was the same for all parameters, with no significant difference between the slopes for the two vegetation conditions in all four dance parameters. Additionally, there was no effect of distance on the slope in the case of the number of dance circuits also. Thus, the presence of intermediate dances does not affect the results obtained in our main analysis.

The relationship between circuit duration and distance has been described as monotonic before (Dyer, 2002) and in a previous study two separate linear segments were fit to the non-linear curve linking circuit duration and distance (Dyer and Seeley, 1991). We believe that future studies should take into consideration a non-linear relationship between the waggle phase and distance as the nature of the relationship between the spatial information signal and distance is important to understand the benefits of the signal itself. To this end, we provide a detailed documentation (and accompanying R code) of the non-linear model fitting process employed in this study below.

#### Non-Linear Mixed-Effects Model Analysis

In the case of the three continuous dance parameters that we analysed (waggle phase duration, circuit duration and return phase duration), we had to fit non-linear mixed-effects models. Even though we tried to fit linear mixed-effects models, we could not make inferences from these models, as the model assumptions, particularly the assumption of homoscedasticity of residuals, were violated. On closer inspection of the raw data, we found that the distribution of the three parameters at the level of both, individual runs and mean values per dance, were not normally distributed (e.g., see Fig. 2 and 3).

Fitting simple non-linear mixed effects models in R is straightforward and we describe the process here to make it accessible. We have also attached an accompanying R code that can be used to replicate our model fitting process. We do not go into details regarding the type of non-linear function to fit, as this will depend on the data and the underlying biological phenomenon. One option would also be to fit multiple non-linear functions to the data and choose between these models. Once a non-linear function is chosen, then the next steps are:

1. Obtain reasonable starting values to implement the NLMM
2. Fit the NLMM
3. Perform model diagnostics
4. Obtain the results of the model and plot predicted fits

While steps 2-4 are common to fitting (generalized) linear mixed-effects models ((G)LMMs), step 1 is different and vital for fitting NLMMs. Fortunately the R stats package (installed by default) implements several self-start functions that can be used to obtain starting values (like

SSmicmen() for Michaelis-Menten models, SSlogis() for logistic models etc.). Additionally, several other packages provide self-start functions for models not covered in the stats package. Of note is the aomisc package which provides several self-start functions of biological interest (the following blogpost provides additional details on fitting these self-starting functions: https://www.statforbiology.com/2020/stat_nls_usefulfunctions/). To obtain the starting values (step 1), we fit the simplified non-linear model (without mixed-effects). This can be done either through nls() function in the stats package in case there is only one continuous predictor, or through nlsList() from the nlme package when there are additional categorical predictors (the exact code for obtaining these starting values are provided in the accompanying .R file).

Once starting values are obtained, NLMMs can be fit (step 2) using the nlme() function from the nlme package. The function is implemented differently from how (G)LMMs are implemented in the generally used lmer and glmmTMB packages. The function consists of three parts, specifying the non-linear function, the fixed effects and the random effect separately. The first part, specifying the non-linear function, contains the response variable and how it is linked to the continuous predictor, as well as the parameters associated with the function. In our case, we used a logarithmic regression of the form:

$$Y = a + b * \log(X)$$

Where Y is the response variable (in this case the dance parameter of either waggle phase duration, circuit duration or return phase duration), X is the distance and $a$ and $b$ are the parameters associated with the logarithmic regression. Thus, the first part of the model is written as:

$$(Dance\ Parameter) \sim \mathrm{NLS.logcurve}(distance, a, b)$$

indicating that the non-linear function linking the parameter and distance is NLS.logcurve with the parameters *a* and *b*. Here NLS.logcurve is a convenient self-start function from the package aomisc which is a wrapper for the logarithmic regression model. In the second part of the model, we specify the fixed effects on which *a* and *b* depend. We built two different versions of the model for each parameter to compare the fixed effects:

1. Neither *a* nor *b* depend on the vegetation condition (implemented as a + b ~ 1), representing the situation where the vegetation condition had no effect on the non-linear relationship between the dance parameter and distance
2. Both *a* and *b* depend on the vegetation condition (implemented as a + b ~ condition), representing the situation where the vegetation condition determines the non-linear relationship between the dance parameter and distance

In terms of conventional (G)LMMs, the first and second models are analogous to models in which only distance is a fixed effect and in which distance and vegetation condition has an interaction effect, respectively. We then compared between these 2 models using the AICc value to determine which is the better fitting model for each dance parameter.

With respect to the third part of the model, the random effect, this was the same in both models. We allowed *b* to vary with individual bee ID, thereby implementing a model analogous to random slopes in (G)LMMs.

In addition to this, we provide the starting values for the required parameters in the NLMMs. For the first model, since there was no covariate of condition, only one value each of *a* and *b* needs to be estimated and we provide 2 starting values. In the case of the second model, two values each of *a* and *b* have to be estimated (corresponding to the two vegetation conditions), and hence we provide 4 starting values. We used the default 'nlminb' optimizer with 100 iterations of the optimizer to fit the models.

The model assumptions are verified (step 3) similar to the case of (G)LMMs, and we checked for homoscedasticity and normality of residuals as well as normality of random effects. Since we used the same generic methods, we don't go into detail here (the accompanying .R file has the code we used). In the case of waggle phase duration for cerana, we still faced the issue of heteroscedasticity in the residuals after fitting the NLMM, although it was slightly better than in the case of the LMM. To deal with such cases, the nlme() function implements an optional weights argument to accommodate the within-group heteroscedasticity structure. We used trial and error to determine the best class of variance functions from the available standard classes. We did this by fitting different models each with its own variance function and then visually inspecting the homoscedasticity plot. We found that, for our data, the varPower function, corresponding to power variance function structure with a function coefficient of 10 on the variance covariate *distance*, was able to deal with the heteroscedasticity in the data.

Finally, obtaining the results of the NLMMs, as well as obtaining predictions for plotting are similar to methods used in (G)LMMs implemented via lmer and glmmTMB packages in R. We provide this also in the code, but do not go into details here, as these methods are fairly standard in statistical analysis using mixed-effects models in R.

### Supplementary Tables

#### Table S1

| Species | Condition | Distance | Dances |
|---|---|---|---|
| *Apis florea* | Dense vegetation | 100m | 8 |
| | | 200m | 5 |
| | | 300m | 7 |
| | | 400m | 4 |
| | | 500m | 10 |
| | Sparse vegetation | 100m | 10 |
| | | 200m | 6 |
| | | 300m | 6 |
| | | 400m | 6 |
| | | 500m | 8 |
| *Apis cerana* | Dense vegetation | 100m | 10 |
| | | 200m | 10 |
| | | 300m | 10 |
| | | 400m | 10 |
| | | 500m | 10 |
| | Sparse vegetation | 100m | 10 |
| | | 200m | 10 |
| | | 300m | 10 |
| | | 400m | 10 |
| | | 500m | 10 |

List of dances analysed for each distance for both conditions in both species

**Table S2**

| Parameter | Dense | | | | Sparse | | | |
|---|---|---|---|---|---|---|---|---|
| | Intercept | | Slope | | Intercept | | Slope | |
| | Est (t/z value) | p | Est (t/z value) | p | Est (t/z value) | p | Est (t/z value) | p |
| WP | **-4031.77 (-8.364)** | **<0.001** | **1034.51 (12.006)** | **<0.001** | **1654.76 (2.509)** | **0.015** | **-348.82 (-2.921)** | **0.005** |
| CD | **-9039.96 (-4.679)** | **<0.001** | **2470.37 (7.148)** | **<0.001** | **5657.44 (2.147)** | **0.036** | **-1202.11 (-2.515)** | **0.015** |
| RP | **-4843.12 (-2.546)** | **0.014** | **1407.33 (4.138)** | **<0.001** | 3769.28 (1.449) | 0.153 | -815.33 (-1.73) | 0.089 |
| NC | **2.139 (10.677)** | **<0.001** | 0.0002 (0.289) | 0.772 | 0.488 (1.926) | 0.054 | -0.001 (-1.165) | 0.244 |

Model summary of the mixed-effects models fit for each of the four dance parameters (WP: Waggle Phase duration; CD: Circuit Duration; RP: Return Phase duration; NC: Number of Dance Circuits) in *A. florea*. The estimate (Est) as well as the associated t/z value (t for continuous parameters, z for discrete parameters) for the intercept and slope in both conditions are provided along with the associated p values. The models were implemented with the Dense vegetation condition as the base contrast level, and hence the p value of the slope and the intercept in this case is associated with whether the estimate is different from zero. In the case of Sparse vegetation condition, the estimate value represents the difference in the estimated value between the Dense and Sparse vegetation conditions, and the p value is associated with this difference. Significant differences at the $p < 0.05$ level are highlighted in bold

**Table S3**

| Parameter | Dense | | | | Sparse | | | |
|---|---|---|---|---|---|---|---|---|
| | Intercept | | Slope | | Intercept | | Slope | |
| | Estimate (t/z value) | p | Estimate (t/z value) | p | Estimate (t/z value) | p | Estimate (t/z value) | p |
| WP | **-3675.46 (-10.136)** | **<0.001** | **862.88 (12.099)** | **<0.001** | -515.74 (-1.006) | 0.318 | 134.73 (1.336) | 0.186 |
| CD | **-6266.26 (-5.826)** | **<0.001** | **1768.24 (9.104)** | **<0.001** | 2211.76 (1.399) | 0.166 | -410.93 (-1.438) | 0.155 |
| RP | -1489.78 (-1.562) | 0.123 | **699.94 (4.061)** | **<0.001** | 1389.19 (0.989) | 0.326 | -291.59 (-1.148) | 0.255 |
| NC | **2.249 (12.349)** | **<0.001** | -0.0002 (-0.298) | 0.766 | 0.168 (0.673) | 0.501 | -0.00009 (-0.134) | 0.893 |

Model summary of the mixed-effects models fit for each of the four dance parameters (WP: Waggle Phase duration; CD: Circuit Duration; RP: Return Phase duration; NC: Number of Dance Circuits) in *A. cerana*. The estimate (Est) as well as the associated t/z value (t for continuous parameters, z for discrete parameters) for the intercept and slope in both conditions are provided along with the associated p values. The models were implemented with the Dense vegetation condition as the base contrast level, and hence the p value of the slope and the intercept in this case is associated with whether the estimate is different from zero. In the case of Sparse vegetation condition, the estimate value represents the difference in the estimated value between the Dense and Sparse vegetation conditions, and the p value is associated with this difference. Significant differences at the $p < 0.05$ level are highlighted in bold

**Table S4**

| Parameter | Dense | | | | Sparse | | | |
|---|---|---|---|---|---|---|---|---|
| | Intercept | | Slope | | Intercept | | Slope | |
| | Est (t/z value) | *p* value | Est (t/z value) | *p* value | Est (t/z value) | *p* value | Est (t/z value) | *p* value |
| SD | **155.862 (4.888)** | **<0.001** | **0.530 (5.850)** | **<0.001** | 60.955 (1.411) | 0.164 | **-0.255 (-2.014)** | **0.049** |
| CV | **0.239 (7.919)** | **<0.001** | -0.00016 (-1.956) | 0.056 | **0.093 (2.334)** | **0.023** | -0.00016 (-1.457) | 0.151 |

Results of the mixed-effects models fit for both variation parameters for waggle phase duration in *A. florea*. The estimates, t/z value (depending on whether the distribution is continuous or discrete) and the p value associated with the estimate is provided. The models were implemented with the Dense vegetation condition as the base contrast level, and hence the p value of the slope and the intercept in this case is associated with whether the estimate is different from zero. In the case of Sparse vegetation condition, the estimate value represents the difference in the estimated value between the high and low vegetation conditions, and the p value is associated with this difference. Significant differences at the $p < 0.05$ level are highlighted in bold

**Table S5**

| Parameter | Dense | | | | Sparse | | | |
|---|---|---|---|---|---|---|---|---|
| | Intercept | | Slope | | Intercept | | Slope | |
| | Est (t/z value) | *p* value | Est (t/z value) | *p* value | Est (t/z value) | *p* value | Est (t/z value) | *p* value |
| SD | 17.238 (0.382) | 0.704 | **0.759 (5.692)** | **<0.001** | 85.355 (1.329) | 0.188 | 0.018 (0.092) | 0.927 |
| CV | **0.316 (8.474)** | **<0.001** | **-0.00026 (-2.372)** | **0.020** | 0.027 (0.503) | 0.617 | 0.00003 (0.217) | 0.829 |

Results of the mixed-effects models fit for both variation parameters for waggle phase duration in *A. cerana*. The estimates, t/z value (depending on whether the distribution is continuous or discrete) and the p value associated with the estimate is provided. The models were implemented with the Dense vegetation condition as the base contrast level, and hence the p value of the slope and the intercept in this case is associated with whether the estimate is different from zero. In the case of Sparse vegetation condition, the estimate value represents the difference in the estimated value between the high and low vegetation conditions, and the p value is associated with this difference. Significant differences at the $p < 0.05$ level are highlighted in bold

**Table S6**

| Parameters | Analysis of All Dances | | | Analysis without Intermediate Dances | | |
|---|---|---|---|---|---|---|
| | Dense | Sparse | *p* value | Dense | Sparse | *p* value |
| Waggle Phase Duration | 862.88 | 997.61 | 0.186 | 1075.39 | 863.64 | 0.472 |
| Circuit Duration | 1768.24 | 1357.31 | 0.155 | 1671.92 | 1327.78 | 0.246 |
| Return Phase Duration | 699.94 | 408.35 | 0.255 | 545.13 | 444.73 | 0.704 |
| Number of Waggle Circuits | -0.0002 | -0.0003 | 0.893 | -0.00003 | -0.0010 | 0.285 |

Comparison of the analysis including the whole dataset and the analysis of the dataset without potential intermediate dances in *A. cerana*. The slopes for the Dense and Sparse vegetation conditions, as well as the p value associated with the difference estimate between the slopes is provided

#### Supplementary Figures

##### Figure S1

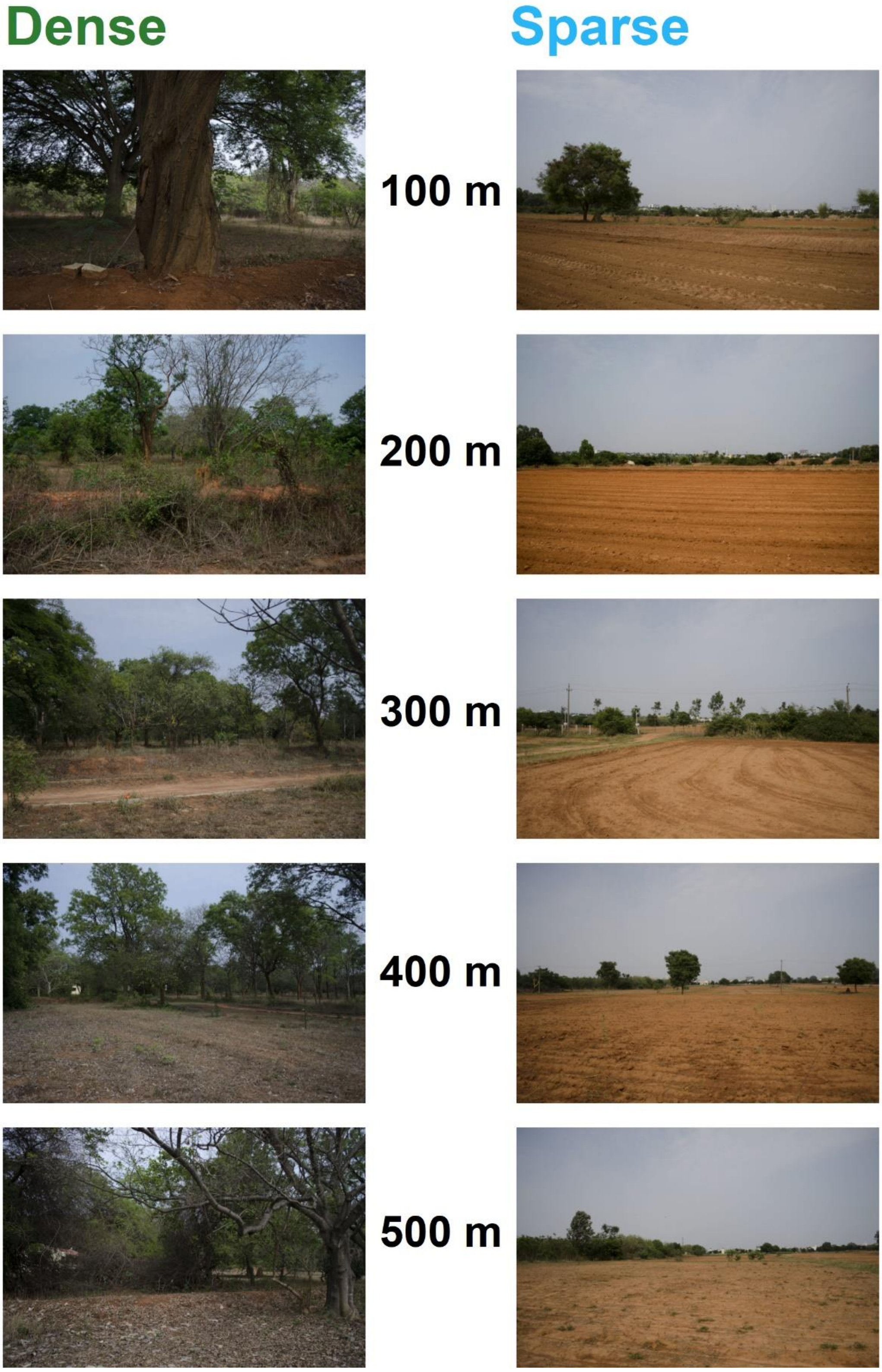


Photographs at every 100 m along the 500 m transect (facing in the direction of the hive) in both the vegetation conditions. These photographs were used to obtain the contrast values depicted in Fig. 1

**Figure S2**

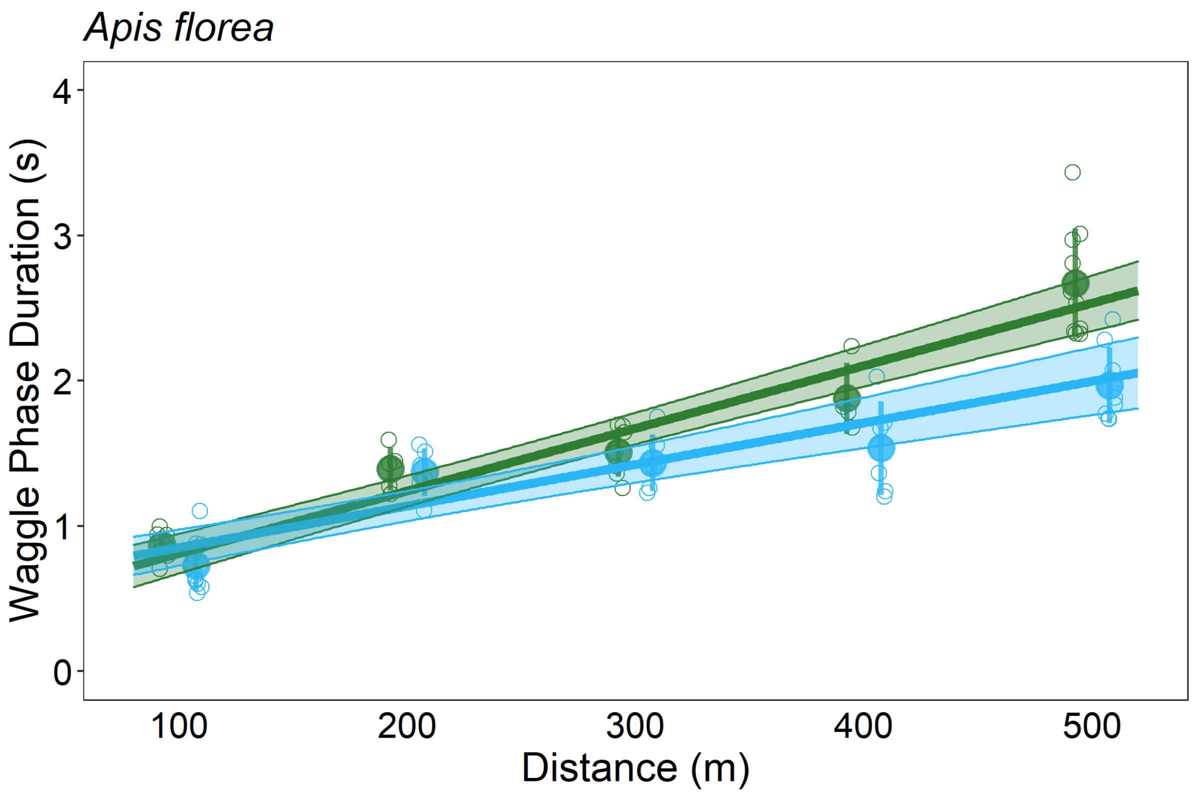


Waggle phase duration with distance in both vegetation conditions in *A. florea*. The mean waggle phase duration for individual dances (open circles), along with the mean of all the dances and the standard deviation (closed circles with error bars) is plotted. Overlaid on top of this is the predicted line and confidence interval associated with this prediction (at the fixed effects level) obtained from the LMM. The circles, lines and confidence interval region are coloured based on the vegetation condition, with green for Dense vegetation and blue for Sparse vegetation.

**Figure S3**

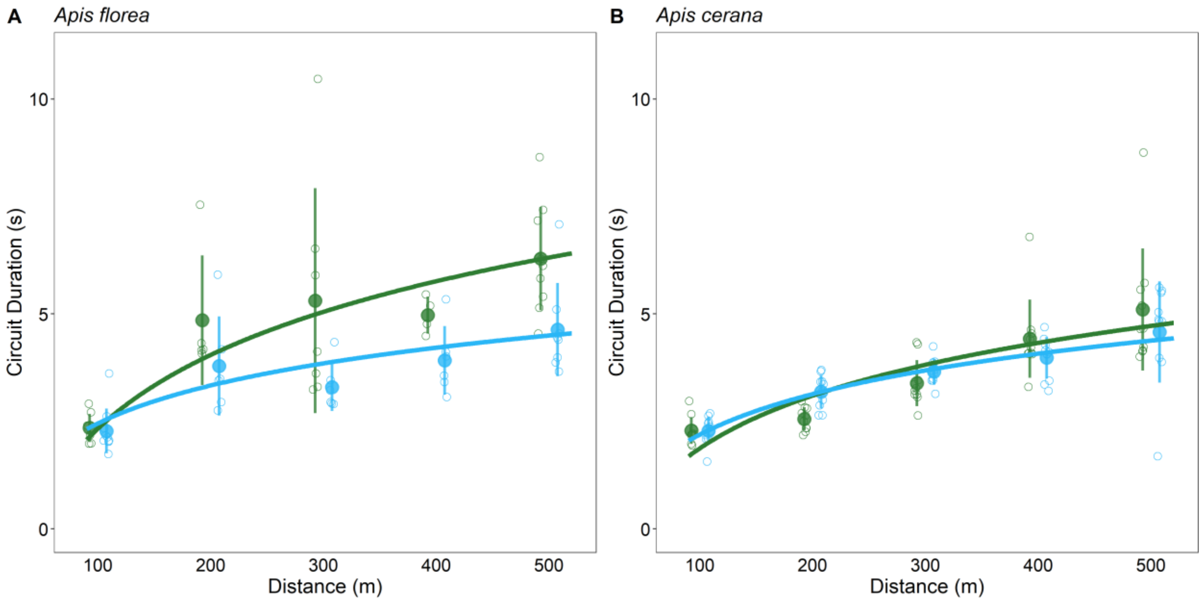


The circuit duration in the two vegetation conditions for A) *A. florea* and B) *A. cerana*. The mean waggle phase duration for individual dances (open circles), along with the mean of all the dances and the standard deviation (closed circles with error bars) is plotted. Overlaid on top of this is the predicted obtained from the NLMM. The circles and lines are coloured based on the vegetation condition, with green for Dense vegetation and blue for Sparse vegetation.

**Figure S4**

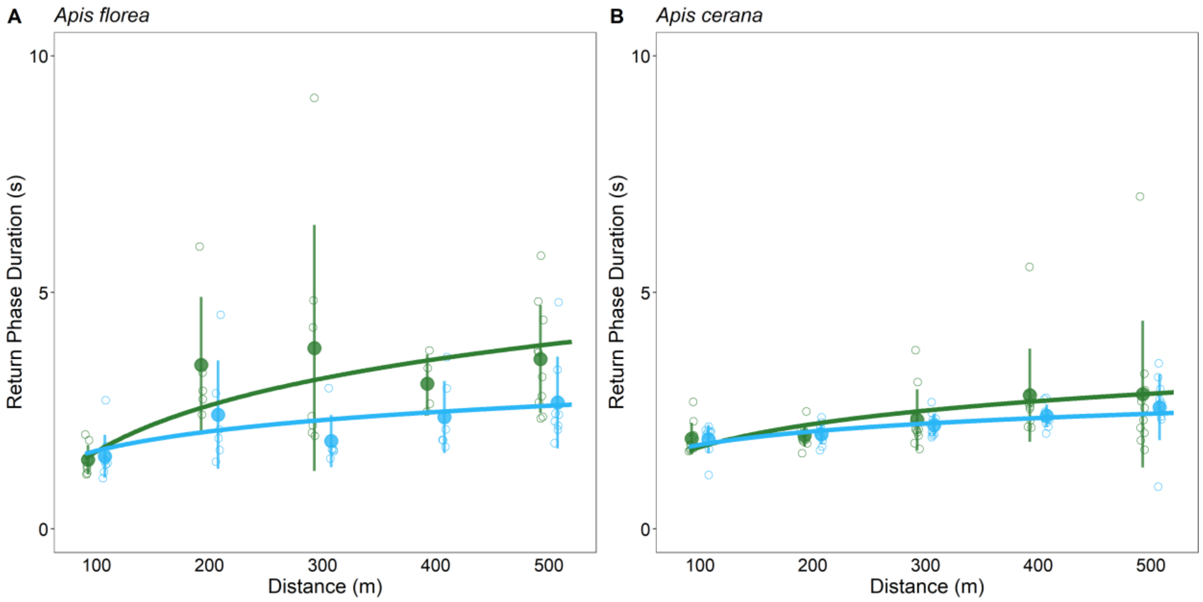


The return phase duration in the two vegetation conditions for A) *A. florea* and B) *A. cerana*. The mean waggle phase duration for individual dances (open circles), along with the mean of all the dances and the standard deviation (closed circles with error bars) is plotted. Overlaid on top of this is the predicted obtained from the NLMM. The circles and lines are coloured based on the vegetation condition, with green for Dense vegetation and blue for Sparse vegetation

**Figure S5**

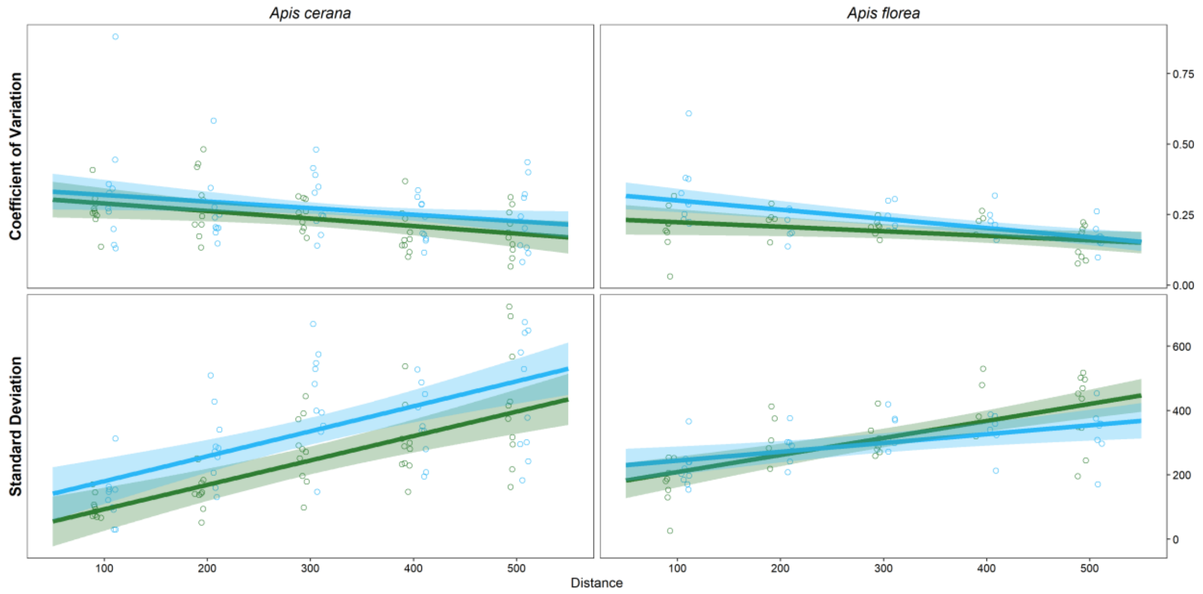


Change in both variation parameters (standard deviation and coefficient of variation) associated with the waggle phase duration with distance and vegetation conditions in both species. Circles represent variation values from each individual dance at each distance. The lines and shaded regions represent the predicted fits obtained from the linear mixed-effects models and the confidence interval (at the fixed effects level) around this fit. The circles and lines are coloured based on the vegetation condition, with green for Dense vegetation and blue for Sparse vegetation
